# Stimulus invariant aspects of the retinal code drive discriminability of natural scenes

**DOI:** 10.1101/2023.08.08.552526

**Authors:** Benjamin D. Hoshal, Caroline M. Holmes, Kyle Bojanek, Jared Salisbury, Michael J. Berry, Olivier Marre, Stephanie E. Palmer

## Abstract

Everything that the brain sees must first be encoded by the retina, which maintains a reliable representation of the visual world in many different, complex natural scenes while also adapting to stimulus changes. This study quantifies whether and how the brain selectively encodes stimulus features about scene identity in complex naturalistic environments. While a wealth of previous work has dug into the static and dynamic features of the population code in retinal ganglion cells, less is known about how populations form both flexible and reliable encoding in natural moving scenes. We record from the larval salamander retina responding to five different natural movies, over many repeats, and use these data to characterize the population code in terms of single-cell fluctuations in rate and pairwise couplings between cells. Decomposing the population code into independent and cell-cell interactions reveals how broad scene structure is encoded in the retinal output. while the single-cell activity adapts to different stimuli, the population structure captured in the sparse, strong couplings is consistent across natural movies as well as synthetic stimuli. We show that these interactions contribute to encoding scene identity. We also demonstrate that this structure likely arises in part from shared bipolar cell input as well as from gap junctions between retinal ganglion cells and amacrine cells.

---

While single cells individually encode specific stimulus features[1–3], it is their aggregate response that drives our perception[4–7]. For this reason, it is important to understand not only how individual cells respond to stimuli, but also how cells influence each other within a population [8–11]. Significant theoretical work has been devoted to understanding population responses [12–16], in tandem with experimental innovations in recording from a large number of cells simultaneously [17–20]. This work has shown that relatively small, diffuse neural correlations are crucial for reproducing the distributions of observed population states or ‘vocabulary’ of the brain. Open questions remain about the functional consequence of these correlations for an organism behaving in its natural environment. This foundation in both theory and experiment on neural population codes creates an opportunity to move to more complex, dynamic stimuli and analyze the code in terms of the readout goals of downstream networks.

The natural environment has many complex spatiotemporal features that make neural encoding in the wild difficult to quantify and assess. Natural scenes vary in luminance over many orders of magnitude [21] and variance [22] [23], and have complicated temporal and spatial structure [24, 25]. Visual systems adapt to these changes on many scales in time and space. Neural systems show near-perfect adaptation to these changes [26], so a question remains about how brains recover scene-specific information once in an adapted state. This open question has led many studies to investigate animal behavior in natural settings [27–30] In this work, we quantify the structure of the neural code at the input end, and how it might support downstream readout that ultimately drives behavior in complex environments.

We use the spike activity in a salamander retina in response to natural stimuli to infer a minimal model of the population structure of its output retinal ganglion cells (RGCs). This structure is, as measured both by statistical predictions and by functional consequences, sparse and conserved across scenes. A human observer of these natural scenes can immediately tell that they are different: how does a small patch of the retina capture scene identity and convey it to the brain? We show that the sparse, consistent correlations between cells helps the population carry long time-scale information about broad scene statistics. Finally, we show that these sparse conserved couplings appear to arise from both shared input (bipolar and amacrine cells) and direct connections (gap junctions).

## Results

### Probing multiple naturalistic, dynamic inputs to the retina

We took dense extracellular recordings from retinal ganglion cells (RGCs) in the larval tiger salamander (Fig. 1a) while presenting 20-second clips from five different movies (Fig. 1b), in a random order, with each shown more than 80 times (Fig. 1c). Salamanders undergo metamorphosis, exposing them to both underwater and terrestrial environments while their retinal structure remains largely the same [31, 32], and navigate through their environment, generating self-motion. The movies chosen represent a sampling of the wide variety of scenes that occur in the organism’s ecological niche (Fig. 1b).

**FIG. 1.**
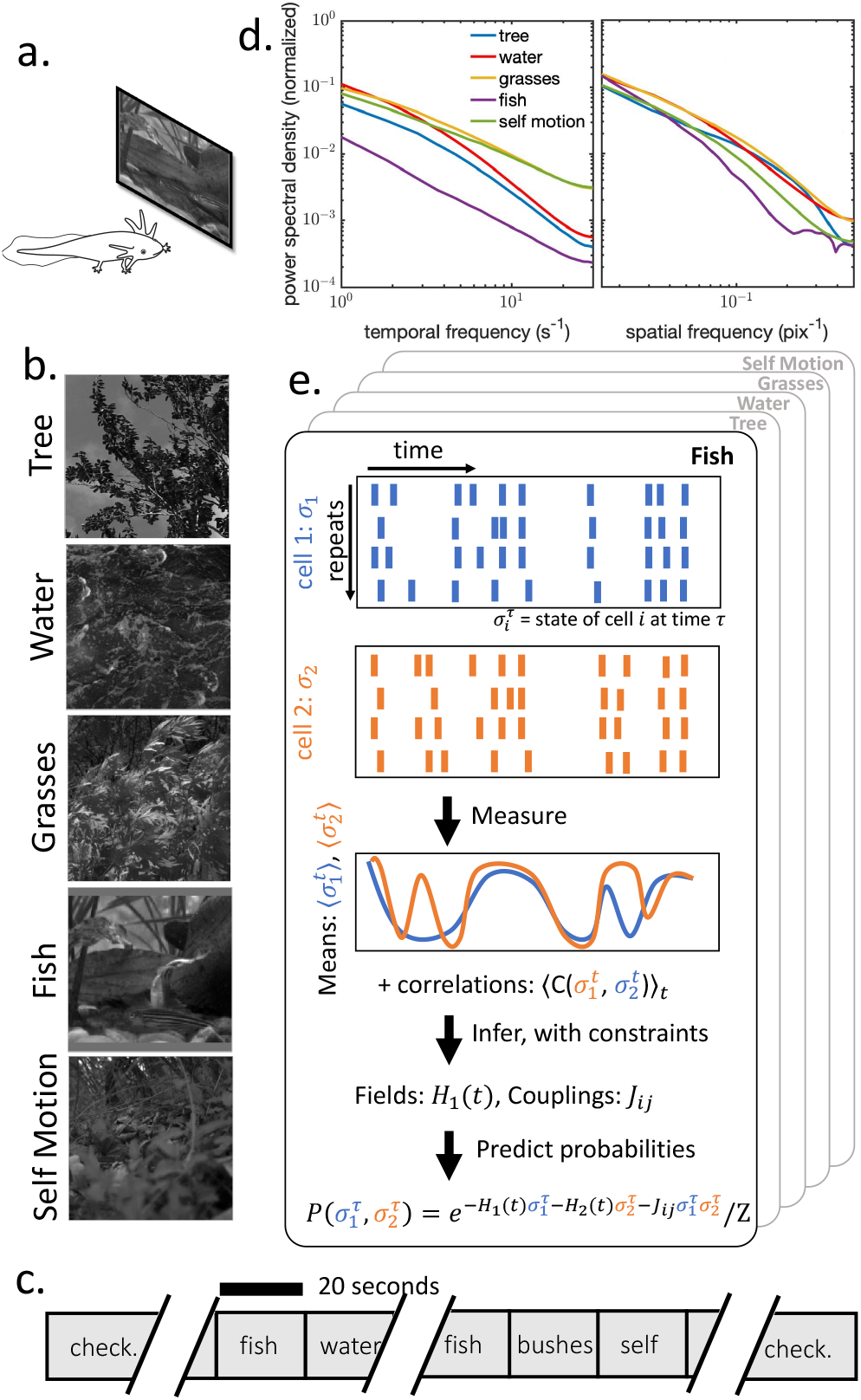
Measuring retinal ganglion cell responses to diverse natural scenes. (a) Voltage responses were recorded from the retinal ganglion cell layer of a salamander retina stimulated by natural movies. (b) Example frames from each of five natural scenes, which show, respectively, trees blowing in the wind; flowing water; ferns and grasses waving back and forth in a breeze; fish swimming; and woodland underbrush as viewed by a moving camera. (c) In order to probe the statistics of responses, natural scenes were repeated a minimum of 80 times in pseudorandom order. A checkerboard stimulus lasting 30 minutes was shown before and after the natural scenes. (d) The five natural movies used have different statistics, including (shown here) spatial and temporal frequency spectra. (e) To model responses to each of the movies, time-dependent maximum entropy models are fit to each of the five natural scenes.

Because of the diversity of subjects and locations, the movies exhibit significantly different temporal and spatial frequency spectra (Fig. 1d). As luminance correlations alone fail to capture behaviorally relevant features like motion, work has quantified the substantially different shapes and timescales of velocity autocorrelations for different scenes [33, 34]. As animals navigate through different environments, they rapidly adapt to these changes in stimulus statistics [26, 35, 36]. Thus, playing a variety of natural scenes in a single experimental session probes a fuller range of neural response repertoires and gives us an opportunity to capture the true range of the brain’s code.

### Pairwise couplings are consistent across movies

To fully describe a dynamic binary population code, we must enumerate 2^*N*^ possible states at each time in the response. Even for modest N, a fully expressive population code is both experimentally inaccessible and potentially unreadable by downstream networks. To summarize the population code, we use maximum entropy modeling, which has a long history of success using *O*(*N* ^2^) parameters to capture the structure in biological data, even recapitulating higher-order features not explicitly constrained by the model [37–44]. We chose these ‘maxent’ models over a Generalized Linear Model (GLM) approach. While GLMs have led to significant insight into adaptation filters in individual cells and effective interactions between cells [10, 45–47], they are not well suited for analyzing the structure of the retinal population code in natural scenes, particularly at the time resolution of interest (16.67 ms, set by the movie frame rate). At this temporal resolution, the primary correlation between cells is the zero-lag correlation, which cannot be modeled by GLMs. In addition, as generative models, GLMs regularly struggle with self-excitation [48].

In applications of maximum entropy techniques in neuroscience, these models are typically constrained by the average firing rate of each cell and the correlation between each pair of cells [37, 49], ⟨σ_*i*_⟩ and ⟨σ_*i*_σ_*j*_⟩. We use a time-dependent maximum entropy model [50] that is constrained by the time-varying firing rates averaged across repeated stimuli. Our model takes the form

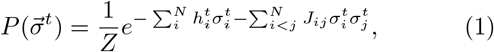

and our constraints are on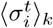, which capture each cell’s individual response to the stimulus at time t averaged over trials, k, (2a) as well as 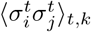,the corre-lations between cells (2b). These two constraints map to two sets of parameters, the time-dependent fields 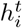 and the static couplings *J*_*ij*_. Interactions between the timedependent fields 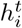 absorb any stimulus-dependent correlations, leaving the couplings *J*_*ij*_ to capture the noise correlations between cells. We independently train mod-els for each movie, fixing couplings as constant across time within movies but allowing them to change across movies (Fig. 1e). This means that couplings are indexed by cell and movie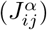, while fields are indexed by cell, movie, and time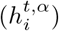.

Our models accurately predict the full suite of popu-lation state probabilities (Fig. 2c), which indicates that higher-order information in the data is well-accounted-for by these second-order constraints. As in other pairwise models fit to neural data, a relatively small number of parameters can capture the full response repertoire of the neural population [37, 51]. This fit is significantly better than the traditional constant-field models without a time-varying field. To quantify this fit, we calculate the overall divergence between the model probabilities and the observed state frequencies, *D*_*JS*_(*P*_*model*_, *P*_*data*_) (Fig. 2d). This analysis again shows that our full model matches our data, while models with constant fields do significantly worse.

**FIG. 2.**
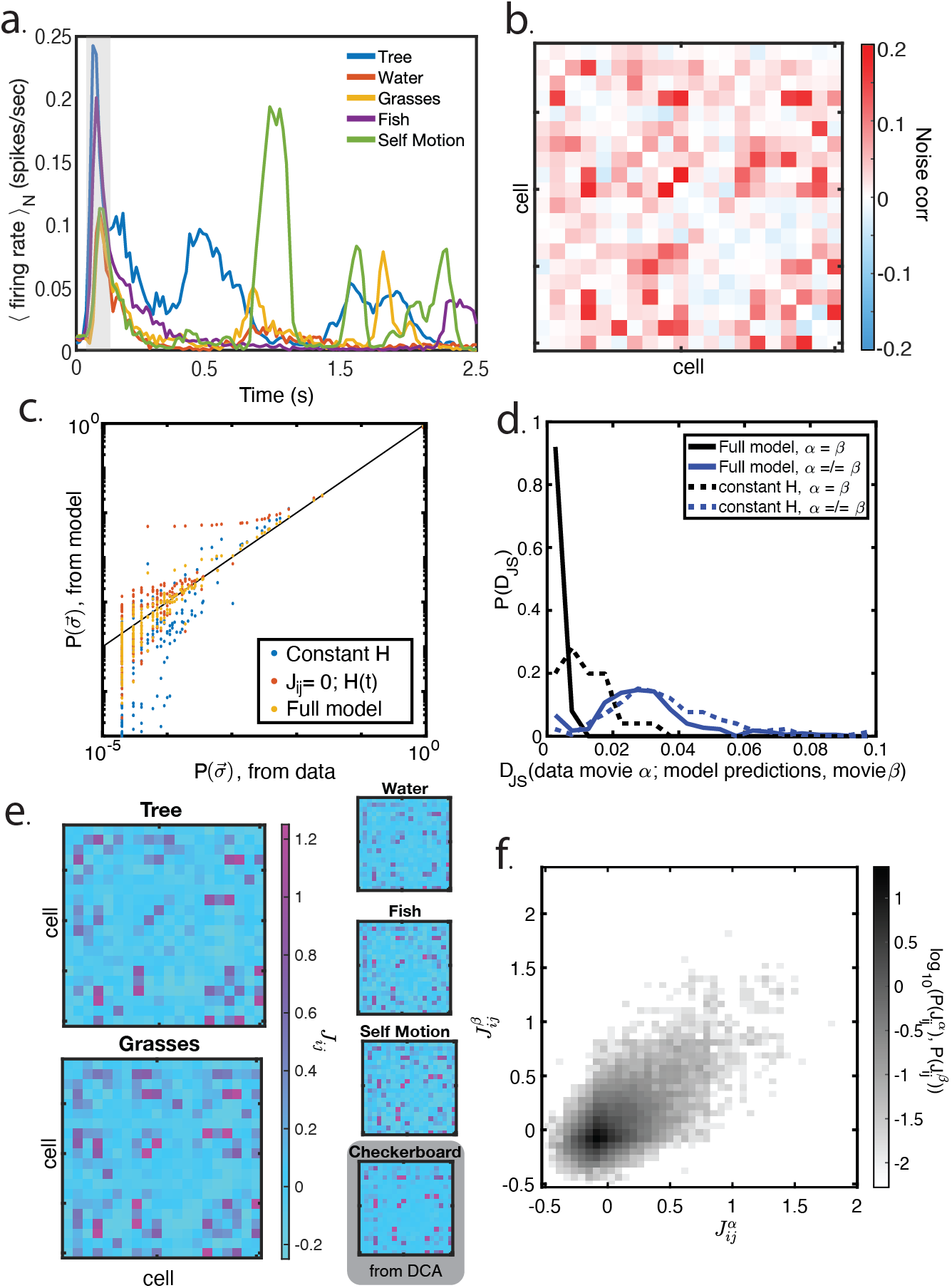
Retinal ganglion cell populations maintain consistent couplings across a variety of natural stimuli. (A)Average population firing rate as a function of time for the first two seconds of each natural movie. In the first 200ms (grey bar), the population firing rate dramatically peaks; after that, rates remain variable in response to stimulus statistics. (B)In addition to time-varying firing rates, cells are characterized by their noise correlations, here shown for a single 20-cell group of cells responding to the ‘Tree’ movie. (c) Probability of population states for a single group of 20 cells, as measured from data and compared to models. The full model successfully extrapolates from second order statistics to population state. Models with either no couplings or with constant fields extrapolate less well. (d) Across a set of ten random choices of 20-cell groups and all movies, our model predicts population states well (low *D*_*JS*_, black line) and the traditional constant-field model does less well (dotted black line). Models across movies are substantially different from each other, and do not predict states well between movies (blue lines). (e) Couplings across all five natural scenes and the checkerboard white noise stimulus, for a single choice of group. In all cases, couplings are consistent across stimuli, especially for the strongest couplings. Even for the checkerboard, where another modeling procedure (DCA) was used to fit the interactions, similar couplings are found. (g) Couplings are quantitatively similar across movies (*R*^2^ = 0.74), especially for the significantly non-zero couplings. Here, we show the values of the couplings *J*_*ij*_ for all pairs of movies, for ten different groups of 20 cells each.

Given the quality of predictions from our models, and that the fields are explicitly constrained by stimulus-induced single-cell statistics, we assume that the interaction matrices *J*_*ij*_ carry the essence of the intrinsic population structure, while the fields carry information about independent responses to the external visual stimulus. We find that these *J*_*ij*_ matrices appear to have similar structure across movies (Fig. 2e). The consistent couplings are not unique to the particular 20-cell group analyzed in Fig. 2e. For a selection of randomly chosen groups, we plot the coupling 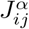 between cells *i* and *J* in movie α against the couplings 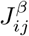 We observe a strong correlation between the sparse set of large-magnitude cell-cell interactions from movie to movie (Fig. 2f).

Previous work has similarly found consistent cell-cell couplings or cell-cell noise correlations across visual inputs [50, 52, 53]. Our goal is to dissect this sparse, consistent ‘skeleton’ of interactions more fully and explore both the origins and functional consequences of this structure. This is the first time that the correlation between couplings has been demonstrated across a range of naturalistic, dynamic stimuli. This consistency is particularly surprising given that these are independently trained models, which separately learned similar couplings despite substantially different scene statistics (Fig. 1d) and predicted population activity (Fig. 2d). This implies that this structure arises from the retina itself, rather than being inherited from the stimulus.

To test whether the similar couplings might arise due to shared long-timescale and length-scale correlations in natural scenes [24], or if they are a property of the feedforward, functional anatomy of the retina, we compare the couplings fit in the natural movies to those found in response to an entirely different, non-naturalistic stimulus.

The population response to a white-noise checkerboard stimulus also has the same kind of pairwise interactions we inferred in resposne to the natural movies (Fig. 2)e. Due to a lack of repeats of the white noise stimulus, we used a different method, Direct Coupling Analysis (DCA) [54], to infer these couplings (see Methods for details). Despite the many differences between this model and those fit to the natural movies (both stimulus and model details), we extract the same strong coupling structure. This strengthens the argument that the observed couplings are indicative of real biological interactions, not correlations inherited from the input.

### Sparse, consistent couplings capture population response structure

The distribution of values in the *J*_*ij*_ *matrices* is heavy-tailed (Fig. 3a, inset), which means that there are rare, sparse strong values. This sparsity of correlated, strong couplings connects with previous work that has highlighted the possible functional role of heavy-tailed coupling distributions in neural networks [55]. Ideas from sparse coding theory have permeated discussions of how large neural populations both store and retrieve information (ganguli2012compressed, ebrahimi2022emergent, candes2006stable). We will quantify this sparsity and consistency, and investigate their role in determining the retinal population code.

**FIG. 3.**
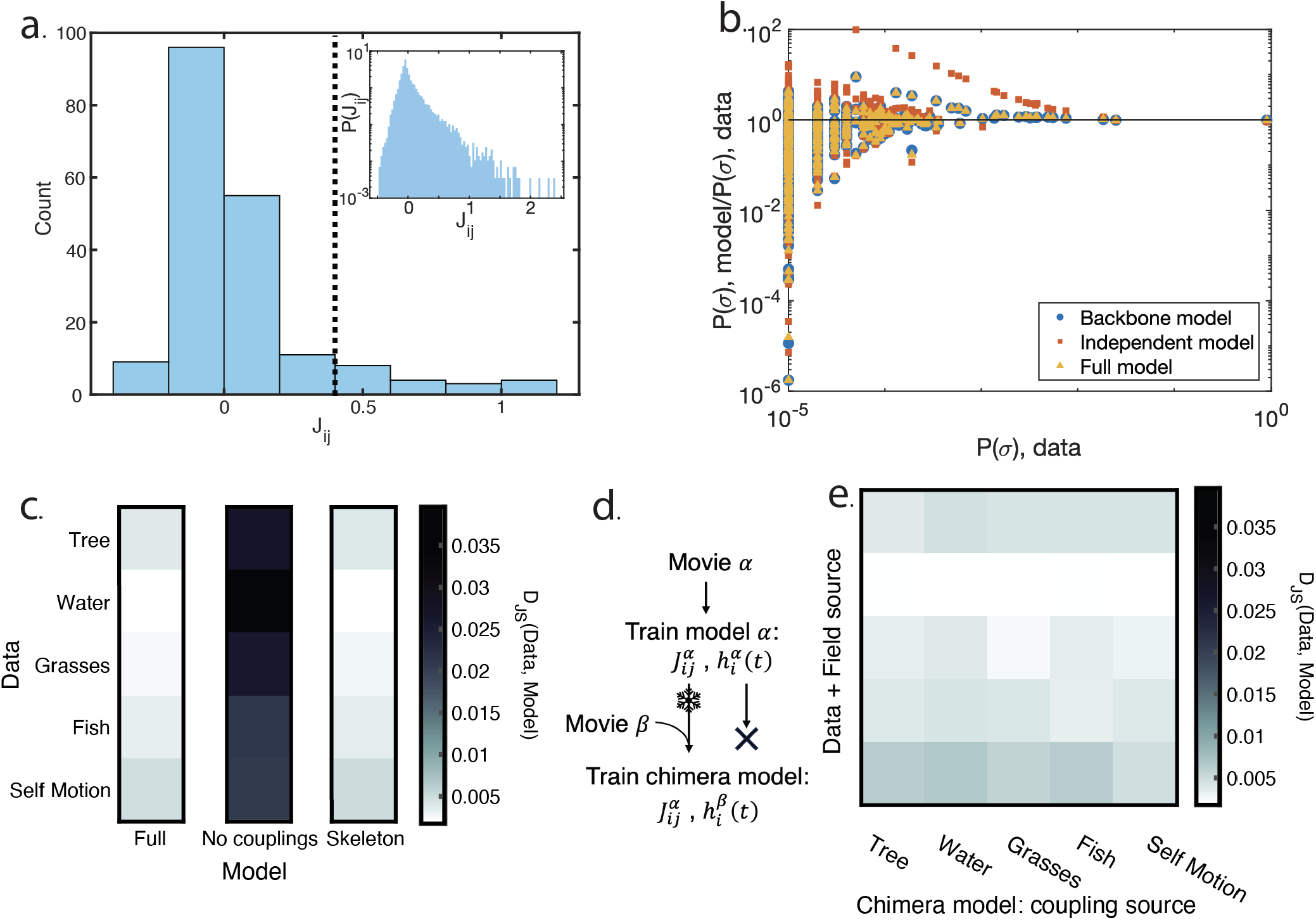
The conserved, sparse coupling structure captures population activity. (a) The distribution of *J*_*ij*_ s is heavy-tailed (inset), and in any particular 20-cell group there are a small number of large couplings. We study this by choosing the top 10% of couplings (above the dotted line), which we will refer to as the skeleton. In general these couplings can be either positive or negative.(b) Probabilities of population states, for our full model, an independent model (all *J*_*ij*_ = 0), and a model trained with only the skeleton values allowed to be nonzero. The full and skeleton models perform nearly identically. (c) Across movies and 10 different groups of 20 cells, the skeleton model performs as well as the full model (i.e., the left column is similar to the right column). (d) To investigate the effects of the variation in couplings, we define ‘chimera’ models, where the couplings are learned from one movie, and then the fields are trained from a second movie. (e) The chimera models perform nearly as well as the full model (each column is nearly identical to the full model column in panel c). We note that the diagonal entries of this matrix are exactly the values in the full model column of panel c.

One measure of the effective sparsity of the *J*_*ij*_ matrices is to check the role of the small set of large couplings in determining population states. Specifically, we set 90% of the smaller magnitude couplings (both positive and negative) to zero. We then refit the remaining, strongest 10% of the *J*_*ij*_s (an example is shown in Fig. 3a). The resulting skeleton model must deviate from the full model in its fit to the true population response probabilities, but it does significantly better than a conditionally-independent model with no *J*’s at all (Fig. 3b for an example of one 20 cell group).

We can broaden this analysis to look at the *D*_*JS*_ between the data and our model-predicted probability distributions, averaged over the choice of 20-cell group (Fig. 3c). We find that the *D*_*JS*_ is small between the data and the full model (as previously shown in Fig. 2d), and increases significantly for a model without couplings. The *D*_JS_ for the skeleton model is nearly identical to that of the full model, meaning that this sparse set of couplings captures almost all of the population response structure embedded in the full *J* matrix.

We can use this analysis to investigate the role of the *changes* in the couplings on the predicted response distributions. We do this by making chimera models, where 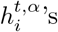 are fit as in the full model, but couplings are inher-ited from a different movie,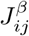 (Fig. 3d). If the model with fields 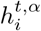 and 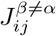 makes very different predictions from a model with 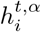 and 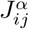, it means that the *J*’s specific to that movie matter. Instead, we find that models with the same α and various β choices perform similarly (evidenced by the horizontal bands in the right part of Fig. 3e). The differences in coupling have minimal effects on determining population activity. Importantly, this is not a statement that couplings do not matter - instead, the *variation* in the couplings has minimal effect, and that they seem to be consistent across scenes.

### Sparse, consistent couplings allow for faster decoding of scene identity

While the results in Fig. 2 show that couplings are consistent across movies, it is not clear from that analysis alone what this means for downstream readout. In general, while couplings between cells have been inferred and their structure discussed in a wide range of neural populations [37, 51, 56, 57], the functional consequence of these couplings is less explored. The couplings observed in these data are both consistent across natural scenes and heavy-tailed (Fig. 3a). Previous work has shown that weak couplings combine to have a large effect on population activity [37], while others have suggested that ignoring weak interactions has minimal effect on population responses [40, 42]. Exploring a broad set of visual scenes allows us to tease apart these two seemingly competing claims about the functional consequence of cell-cell couplings in the retina.

While our preceding analysis shows that the core predictive power of our coupling models are invariant to elimination of all but the largest 10% of interactions, and that the changes in coupling observed across movies do not significantly perturb model predictions, the real biological system might *not* care about inferring the population state distribution. A more compelling test of the role of couplings would be to investigate their role in a downstream behavioral or cognitive task.

Previous work has investigated the effect of interactions between retinal cells on stimulus encoding and retinal development [8, 12, 44, 58–60]. In contrast with this work, we focus on the decoding problem of inferring scene identity. The sparse, strong couplings may combine with the time-dependent fields to signal broad scene spatiotemporal structure. The movies contain a suite of higher order features that make each one readily discriminable to the human eye, but may also impact the local, correlated retinal population code in a readable way.

To test whether the couplings affect the discrimination of scenes, we take advantage of the fact that each of our movies comes from a significantly different environment, which means that the information about scene statistics can be approximated by information about scene identity (1d). We can then quantify the accumulation of evidence about scene identity in a Bayesian framework, given an ideal decoder with the posterior *P*(movie|spikes). This decoding task is similar to the real problem solved by downstream brain areas when an organism moves between scenes as it navigates its natural environment. The brain must invoke different behavioral repertoires and priors in different scenes.

We use a Bayesian approach to measure *P*(movie|spikes),

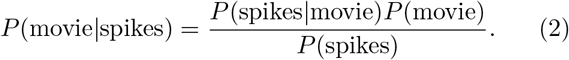

In order to calculate this quantity, we sample population activity from each model that we fit, either making sparse *J*’s, eliminating couplings entirely, or mixing couplings between scenes (see Methods for details). Importantly, because we generate data from reduced or mixture models at will, we can quantify how much each component of the model matters for inference of scene identity.

We find that samples generated from even a 20-cell group carry enough information to correctly identify a scene within a few seconds (Fig. 4b, solid black line). With more neurons, even faster decoding is likely possible. By contrast, spikes generated from a conditionally independent model, with all *J*’s set to zero, take nearly twice as long to achieve the same confidence level in scene identity. We emphasize that the only difference between these models is the couplings; both models have timevarying fields, which capture the spatial and temporal correlations in the natural scene. The disparity in performance is surprising, as it was not *a priori* obvious that couplings would or should contribute to decoding [61]. Prior efforts to establish the functional role of couplings, especially in natural scenes, have shown that correlations matter for discriminating between spontaneous and stimulus-driven responses [14]. To push this further, we can examine how couplings support discrimination *between* complex stimulus-driven ensembles.

**FIG. 4.**
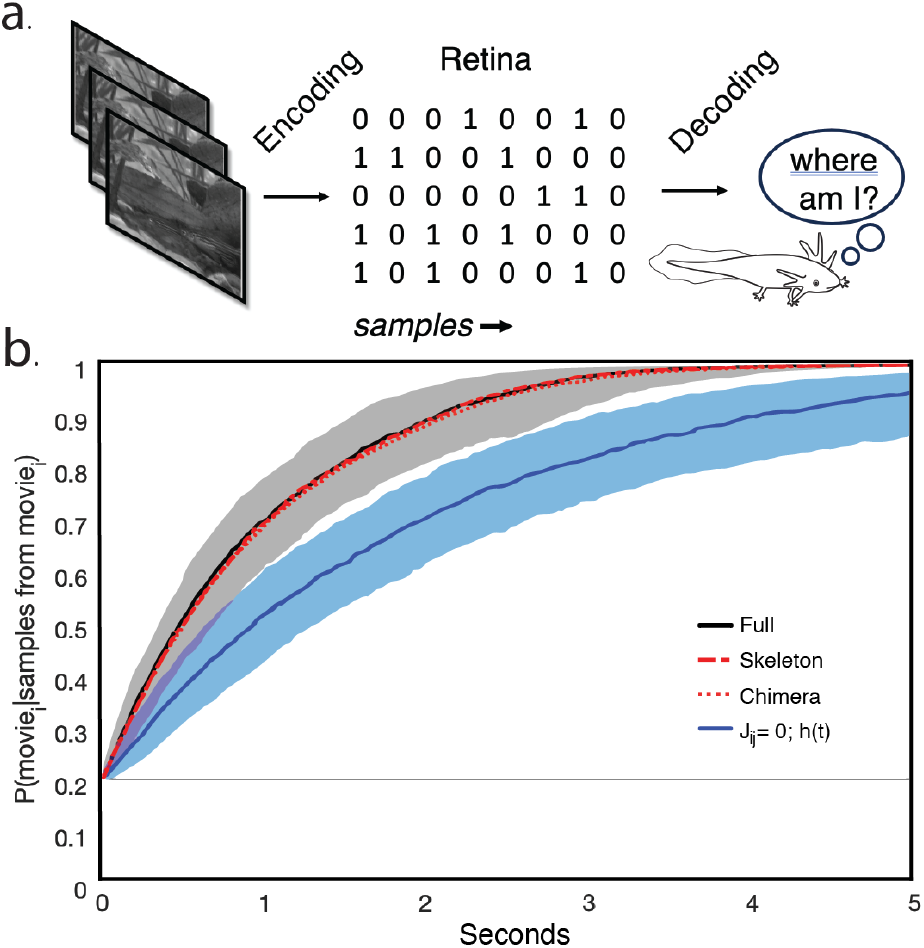
The conserved, sparse coupling structure contributes to readout of scene identity. (a) Cartoon of how we consider the decoding problem downstream of the retina. Signals pass to the retina and are encoded, they then must be decoded downstream. Here, we will consider the task of decoding scene identity as a proxy for decoding large-scale scene statistics. (b) Decoding performance of a variety of models, as measured in number of seconds to achieve high certainty about the correct scene identity. There is a significant difference between the performance of networks with or without couplings, but no difference between networks with the full set of couplings, with only the skeleton couplings, and networks with swapped couplings.

A sparse coupling matrix also supports fast inference of scene identity. We calculate the accumulation of evidence from samples from a skeleton model, where only 10% of couplings are nonzero. Evidence about scene identity accumulates just as rapidly as that of the full model (Fig. 4b, dot-dash red line). The benefit to decoding that arises from the couplings is solely contained in the few strongest couplings, and weaker couplings do not play a measurable role in reading out scene identity.

We additionally investigate the functional effects of the variation in couplings across movies. To do this, we calculate how quickly evidence accumulates for the correct scene in the counter-factual case where the couplings between movies have been swapped between scenes. In that chimera case, we again find that the performance is identical to that of the full model (Fig. 4b, dotted red line).

### Couplings arise from both gap junctions and shared bipolar cell input

We have found consistent population structure that is dominated by sparse couplings. This structure is hard-wired into the retina code and could arise from many different local circuit mo-tifs. Pairwise retinal couplings could be correlated with shared upstream input from a bipolar cell (Fig. 5a, left), direct gap junctions between retinal ganglion cells (Fig. 5a top), or gap junctions between RGCs and a third neuron such as an amacrine cell (Fig. 5a right). All of these mechanisms have been previously implicated in phenomena that could play a significant role in computation in natural scenes. Lateral connectivity (through amacrine cells or gap junctions) is involved in many anticipatory behaviors in the retina [62–64], while feedforward input from bipolar cells to RGCs is involved in many adaptation phenomena [65, 66].

**FIG. 5.**
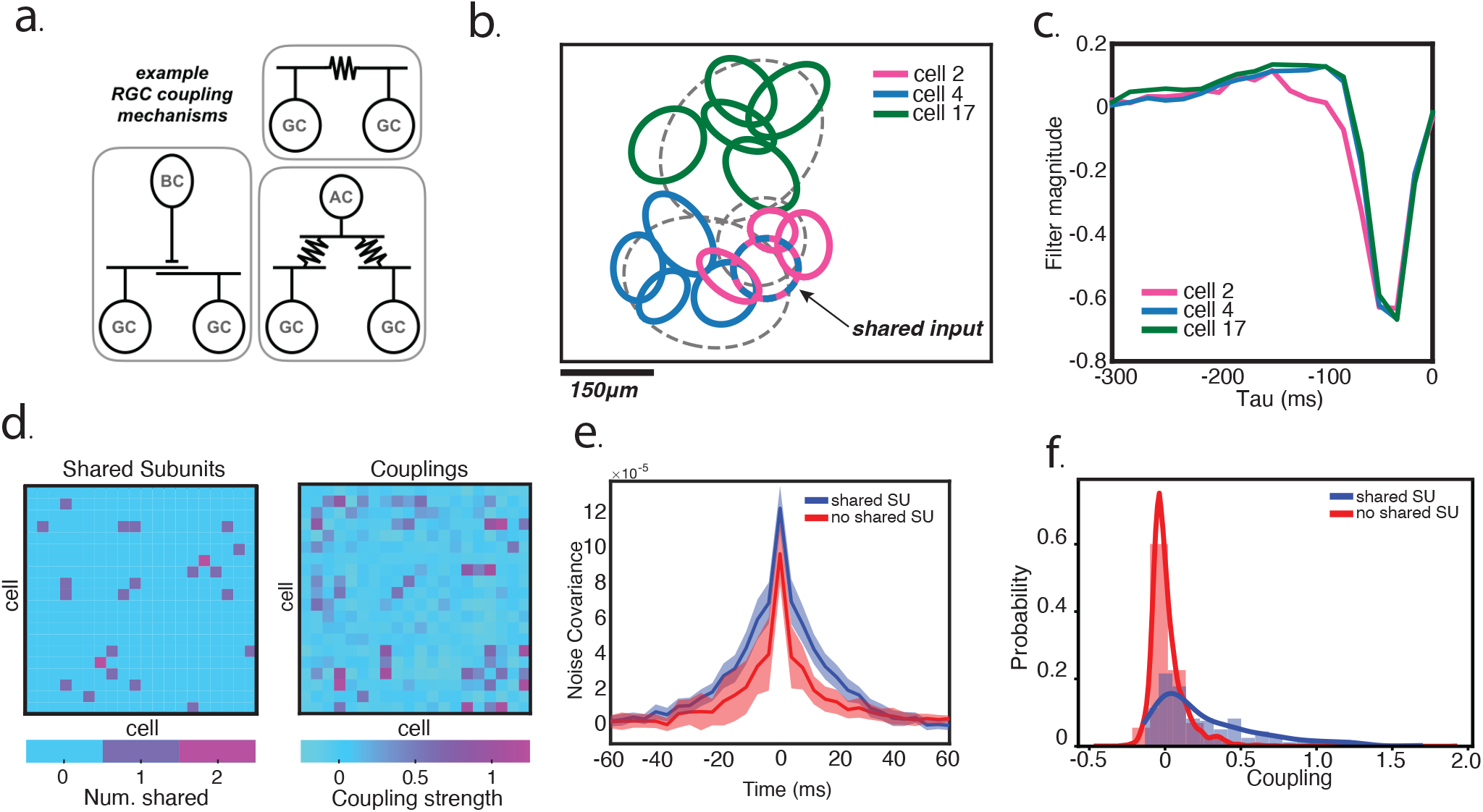
Strong couplings result from both shared inputs and gap junctions. (a) Three potential pairwise coupling sources. Left, shared input from an upstream bipolar cell. Top, a direct gap junction between two retinal ganglion cells. Bottom right, gap junction connections to a shared third neuron, an amacrine cell. (b) ST-ICA modeled spatial subunits for three cells, two of whom share a subunit. (c) Time filters for the same cells shown in b. (d) A comparison between shared subunits and couplings indicates many highly coupled cells shared upstream input. (e) The average noise covariance for highly coupled cells. Highly coupled cells that share a subunit have a broader noise covariance in time than coupled cells without a shared subunit. (f) Distribution of coupling strengths, both for cells sharing subunits and those that do not.

Nonlinear summation of bipolar cell (BC) inputs has been shown to be an integral component of retinal computation [67–72]. The convergence of BCs onto RGCs is modeled using nonlinear summation in so-called cascade models. These models explain a wide array of complex retinal computations (e.g. motion onset [67], omitted stimulus response [73], background vs object motion [74], reversal response [75–77]). As a complement to that, BCs have diverging projections onto multiple RGCs on the retina [78]. While bipolar cells sample a smaller portion of the visual scene than RGCs, gap junctions networks between RGCs can expand the bipolar projective field out as far as ∼1mm [79]. This BC divergence could play a role in shaping the population code in the retina and needs to be further explored with naturalistic, dynamic inputs.

In order to detect putative bipolar cell inputs onto the RGCs in our dataset, we use spike triggered independent component analysis (ST-ICA) [80] on the white noise checkerboard stimulus. ST-ICA models each RGC as the output of a temporal filter and spatial subunits. Similarly, these subunits have been experimentally shown to map to bipolar cell inputs [81]. We show an example of the spatial subunits and temporal filters for three fast-OFF RGCs in Fig. 5b and Fig. 5c, two pairs of which (blue, pink; green, pink) demonstrate strong couplings. One coupling pair (blue,pink) exhibits a highly overlapped spatial subunit that we classify as a shared upstream input (see Methods). In Fig. 5c, temporal filters for two cells (blue, green) show nearly identical characteristics. This may arise from a gap junction connecting them and might explain their strong coupling [50].

Across the population, many RGCs share spatial subunits. These subunits closely align with the observed coupling matrix from the stimulus dependent maximum entropy models (Fig. 5d), demonstrating that strong couplings may arise in part from shared upstream input. However, not all cells with strong couplings share upstream input, and the presence of a shared subunit alone does not guarantee the existence of a strong coupling. Thus, strong couplings might arise from multiple sources within the retinal cell population.

Previous work has suggested that gap junctions may underpin couplings between RGCs [8, 50]. Here, we find evidence of gap junctions by inspecting the cross-covariance of responses after subtracting the trial-average (at zero lag this is the usual noise correlation). Pairwise noise correlations due to gap junctions can generally be split into two classes, direct RGC-RGC couplings and shared gap junctions with a third upstream neuron. The symmetric, medium width correlation we observe between some highly coupled cells without a shared subunit (Fig. 5d, red) likely arises from this second class, as direct RGC-RGC gap junctions lead to transient noise correlations at a sub-millsecond timescale [8]. Furthermore, the broader noise correlations between RGCs with shared subunits demonstrates a timescale longer than can be explained from gap junctions alone and may indicate a coupling arising from shared upstream input (Fig. 5a, blue).

Shared input between RGCs, whatever the source, greatly increase the likelihood of a a strong coupling compared to RGC pairs without shared input (Fig. 5e). Of course, shared input does not *guarantee* a strong coupling between RGCs, but the long tail of strong couplings for pairs with shared input suggests that these mechanisms underpin our sparse network of strong interactions. We hypothesize that shared bipolar input and gap junctions work in concert to generate a sparse set of intrinsic correlations between RGCs.

## Discussion

Sparse, strong couplings between cells in this neural population are an important component of both generating the population response repertoire and potential downstream readout of natural scene identity. While some studies show that independent models of the retinal code retain upwards of 90% of the response structure [82], such analyses are typically agnostic to the downstream readout goals of the organism. Without a defined readout goal, it is impossible to determine whether aspects of the response structure lost by an independent encoding relay information meaningful to the organism. It is possible that an independent readout preserves much of the population response range while failing to effectively convey critical features of the visual scene. Natural scenes probe a behaviorally relevant context to assess the impact of noise correlations on neural coding. These movies, like other natural inputs, drive a richer and more reliable code in the brain [83– Comparing across movies reveals what the more subtle features in the neural code might be used for.

Both cell-cell couplings and time-dependent fluctuations in individual cell excitability shape the observed retinal response distributions and convey information about scene features. Previous work has shown that an independent model of retinal activity can recover information about scene features, like spatial texture and local motion ([86]). This study used the trial averaged, time-dependent firing rates across the full retinal population to show how a lower dimensional, latent representation of this activity could support precise readout of time in a natural movie. These latent representations also generalized across different natural movies ([86], in figure 3a). The local time-dependent fields in our maximum entropy model are the key features for this kind of scene encoding, while our sparse skeleton of interactions encodes broader categorical scene information. The interplay between these static and dynamic features provide a rich substrate for encoding different kinds of scene information flexibly and reliably.

Only a sparse subset of strong couplings is critical for scene identity decoding. There is a distinction between this kind of ‘functional sparsity’ (where the couplings matter for performance) and ‘statistical sparsity’ (where the distribution is, itself, sparse). Previous work has shown that cortical populations are both statistically and functionally sparse [87]. This could be a general strategy by which different kinds of information are multiplexed in a neural code [88, 89].

Our finding that sparse interactions sufficiently capture the functional impact of noise correlations on neural encoding elaborates on previous work that argued for a dense network of weak couplings in the retinal code [37, 41]. In these early models, both the fields and interactions between cells were static. The fluctuating fields we included (following the time-dependent maximum entropy model proposed in [50]) absorb much of that structure, and capture the independent component of stimulus-driven changes in the neural population response. The remaining sparse couplings are potentially the key factor for efficient scene identification. A sparse skeleton of interactions may be flexibly activated in different scenes in a way that is easier to discriminate downstream. These interactions generate noise correlations that may reflect changes in external scene correlation structure, which may help recover scene-specific information that is otherwise lost to single-cell-level adaptation [26]. Notably, these interactions allow for scene decoding at a similar timescale as some behavioral features in amphibians [90–92]

On the flip side, sparse codes might hamstring error correction [93, 94], so future work should explore how these costs and benefits trade-off for behaviorally relevant inputs and tasks. New holography work may provide the kind of detailed probe needed to explore the dual roles of individual cell responses and cell-cell correlations in natural tasks [95–97].

These data provide some hints at the mechanisms for generating this sparse but strong functional connectivity in the retina. In many ways, the circuit structure in the eye differs from that found in the cortex, yet some common anatomic motifs are found in both. The retina is not a recurrent neural network, RGCs do not have direct synaptic coupling, and the photoreceptor-to-RGC circuit is largely feed-forward. However, gap junction coupling is prevalent throughout the brain, as is shared, convergent input to regional projection neurons. To create a population code with sparse interactions, the retina needs to be wired around its particular structural constraints, but appears to use these common motifs found throughout the brain. Our initial results indicate that sparse interactions may be the result of common bipolar inputs and gap junction coupling between RGCs. Both gap junctions and common bipolar inputs lead to stronger coupling between cells, but our analysis is not sensitive enough to tease apart whether these two types of coupling sources are mutually exclusive in particular pairs of cells. Exclusivity would be an efficient way to implement a sparse skeleton of specific cell-cell interactions. Future work to disentangle the circuit mechanisms giving rise to the sparse skeleton might ultimately inform studies in cortex where gap junction coupling is also present [98–100].

## Methods

### Neural data

Voltage traces from the output, retinal ganglion cell layer of a larval tiger salamander retina were recorded following the methods outlined in [17]. In brief, the retina was isolated in darkness and pressed against a 252 channel multielectrode array. Voltage recordings were taken during stimulus presentation of both natural movies and white noise stimuli, and spike-sorted using an automated clustering algorithm that was hand curated after initial template clustering and fits. This technique captured a highly overlapping neural population of 93 cells that fully tiled the recorded region of visual space. Spike times were binned at 16.667ms for all analyses presented.

### Visual stimuli

A white noise checkerboard stimulus (with binary white and black squares) was played at 30 frames per second (fps) for 30 minutes prior to and after the natural scene stimuli. Five different natural movies lasting 20s were played in a pseudorandom order, and each were displayed a minimum of 80 times. The movies labeled tree, water, grasses, fish, and self motion were repeated 83, 80, 84, 91, and 85 times, respectively. All natural scenes except for the tree stimulus were displayed at 60fps. The tree stimulus was updated at a rate of 30fps with each frame repeated twice to match the 60fps frame rate of the other movies.

In all movies, the cells significantly increase their firing rates in the first 200 ms (Fig. 2a) following the switch to a new stimulus. This is followed by a rapid decay back to a baseline firing rate. This is likely due to a strong population response to abrupt changes in luminance within their receptive fields [65, 101, 102]. In subsequent analysis, we exclude the first 500ms of every trial to isolate the more steady-state response of the retina to scene-specific features and dynamics.

### Maximum entropy modeling

We followed the datadriven algorithm introduced in [103] for our maximum entropy modeling. This algorithm uses a quasi-Newtonian method that allows for inference of model parameters **X**, in our case time dependent fields 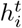 and constant couplings *J*_*ij*_, without needing to compute the inverse model susceptibility matrix χ^−1^[**X**] at each time point during the learning dynamics. As in [50] we learn a maximum entropy model with time-varying fields. Specifically, we learn a model of form

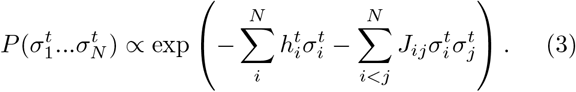

The time dependent fields 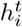 capture the time-varying firing rate of cells σ_*i*_ in response to the stimulus. This means that stimulus dependent correlations between cells are absorbed into the fields, leaving the couplings *J*_*ij*_ to encapsulate the noise correlations between cells.

All fits were done on groups of 20 cells, which were all subsets of the full population of 93 cells. These subsets were chosen at random.

We additionally used data from sparse models to generate the skeleton and independent information shown in Fig. 3. In order to train the independent model, we follow the same procedure as our full pairwise models, but we constrain all couplings to be zero at all fitting steps. In order to train the skeleton model, we first fit a full model with all couplings and fields. We then select the top 10% of couplings, and refit a model where the only allowed nonzero couplings are pairs identified in those top 10% of couplings.

Model performance was tested by comparing predicted probabilities of the population response states to observed frequencies in the data. This is a dramatic increase in the number of parameters from *N*^2^ to 2^*N*^, and represents a difficult extrapolation task.

### DCA models

When we were looking for a model that would give us access to the couplings in data from the white noise stimulus, we were faced with the lack of stimulus repeats. The stimulus-dependant maximum entropy model fundamentally relies on measurements of noise correlations, and as such does not work without stimulus repeats. However, we also wished for a method that would allow us to infer couplings that are only representative of the noise correlations, not couplings that included both shared stimulus inputs and noise correlations. In general, this is not possible, and standard maximum entropy model are not up to the task. Here, however, we took advantage of a feature of these data: many of our cells have highly overlapping receptive fields. This means that two cells with similar spatial receptive fields can be correlated two ways: because of a noise correlation, or because of shared stimulus drive. In particular, if several cells are driven in the same way by the stimulus, they will all have correlations with each other, and therefore many methods would infer ‘loops’ of couplings.

However, in the protein community, DCA (Direct Coupling Analysis) was developed as a maximum-entropy technique with an emphasis on ignoring indirect couplings (i.e., breaking loops) [54, 104], and a prior on the sparsity of the coupling matrix. This means that in our case, with several cells all driven locally in the same way by the stimulus, many of those correlations will be dropped in favor of a sparser explanation for the population activity. In particular, one should expect the remaining couplings to be the strongest correlations - those where there is both a biological coupling between the cells and a shared receptive field. This means that we do not expect quantitative agreement between this method and stimulus-dependant maximum entropy, but that we can hope to find the same skeleton of strong couplings.

In order to fit the DCA model, we followed methods discussed in [54]. We chose a gauge where neural silence or activity are described by {0,1} to more easily relate the DCA network to couplings fit from the time-dependent maxent models.

### Decoding scene identity

For the scene identity decoding introduced in Fig. 3b, we used a Bayesian approach to measure *P*(movie|spikes)

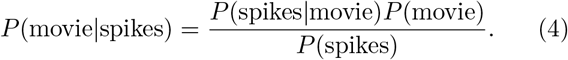

This describes the decoding task - deciding which of the five movies is being shown, based on the population activity from 20 neurons. This is highly related to the problem solved by downstream networks reading out retinal output and attempting to determine large-scale information about the environment. In this case, the decision is between five equi-probable scenes, so we have a uniform prior, *P*(movie) = 1/5.

In order to calculate *P*(spikes|movie), we need to sample spike trains from our models. Our goal is to evaluate the ability of various underlying generative architectures to support scene discriminability. Sampling from the models yields two key advantages: first, we can generate as many samples as needed from those distributions, and second, we can directly investigate the effects of degrading couplings or swapping couplings between scenes. ‘Spike data’ for the model will be a set of vectors 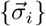, where each is the binary representation of spiking of all neurons in time bin *i*. The *J*’th element of 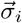 represents neuron *J*’s activity in time bin *i*. All of our models produce a time-dependent probability distribution of joint activity of the neurons, 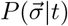, but also produces a timeindependent distribution, 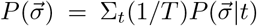. We will generate each spike vector 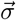 by pulling from this time-independent distribution. We do this separately for each time bin in our simulated spike train.

Armed with a set of neural responses to a particular movie 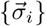, we then can calculate 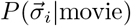 For movie α, we calculate 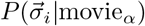 based on each model.

To calculate 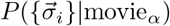 at time point *τ* on the plot in Fig. 4b, we need to accumulate evidence, 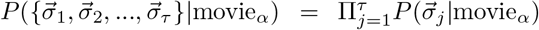 Finally, we can calculate the overall probability of a given spike pattern, 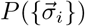, by summing over movies, 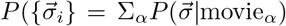 We can easily change the underlying models and resample from the population patterns, allowing for easy comparisons on this decoding task.

In order to generate Fig. 4b, for each combination of movie and model choice, we generate 1000 spike trains. We show the median performance of each model at decoding, and we show the quartiles for the full and conditionally-independent models.

### Spike Triggered ICA

To compute the Spike Triggered ICA (ST-ICA) we follow methods developed in [80].

We first compute the spike triggered average for each cell in a natural cubic spline basis. This is a common method to reduce the number of parameters needed for the model and ensure that the resulting receptive fields are smooth in space and time. We choose the number of splines such that the log-likelihood on held out white noise data is maximized.

To further reduce the number of parameters, we assume our receptive field is rank 1, it can be separated into a spatial filter and a temporal filter. Following these assumptions we use SVD to find this rank 1 approximation. This provably minimizes reconstruction error under the Frobenius norm. We then crop the spatial dimensions for each cell to the regions containing the receptive fields and convolve the stimulus with the temporal filters, which leaves only spatial degrees of freedom.

For each cell, we then have a matrix of size *N* spikes by *M* features, where each feature is a spatial pixel convolved with time filter. We use Preconditioned ICA [105], an algorithm for ICA that uses preconditioned L-BFGS, a low memory quasi Newton optimization algorithm, for optimization to estimate 20 independent components. Resulting components were considered proper subunit candidates based on the presence of significant spatial autocorrelations, following methods in [81].

With a list of candidate subunits for each cell we then compute the activation of that subunit by projecting the time convolved stimulus onto each filter identified by ST-ICA. Two units are considered the same following methods developed in [106].

## I Acknowledgements

We thank Ulisse Ferrari for helpful discussions and providing a preliminary version of the code used for the time-dependent maximum entropy models. This work was supported in part by the National Science Foundation, through the Center for the Physics of Biological Function (PHY-1734030) and an NSF Graduate Research Fellowship (DGE-2039656) (CMH); and by the National Institutes of Health BRAIN initiative (R01EB026943). This work was additionally supported by the National Science Foundation through the Physics Frontier Center for Living Systems (PHY-2317138), and by the NSF-Simons National Institute for Theory and Mathematics in Biology (NSF DMS-2235451 and Simons Foundation MP-TMPS-00005320).

